# COGENT: evaluating the consistency of gene co-expression networks

**DOI:** 10.1101/2020.06.21.163535

**Authors:** Lyuba V. Bozhilova, Javier Pardo-Diaz, Gesine Reinert, Charlotte M. Deane

## Abstract

Gene co-expression networks can be constructed in multiple different ways, both in the use of different measures of co-expression, and in the thresholds applied to the calculated co-expression values, from any given dataset. It is often not clear which co-expression network construction method should be preferred. COGENT provides a set of tools designed to aid the choice of network construction method without the need for any external validation data.

**Availability and implementation:** https://github.com/lbozhilova/COGENT

## 1 Introduction

Gene expression data is a powerful resource for understanding genetic function under different conditions. A common way of exploring this data is through gene co-expression networks (Lee *et al.*, 2004). In these networks genes are represented by nodes and highly co-expressed gene pairs are connected by edges. Such networks have been used in many ways, including for gene function prediction and the identification of disease- or tissue-relevant gene modules (van Dam *et al.*, 2017).

Gene expression data typically takes the form of a matrix, in which rows correspond to genes and columns correspond to samples. Network construction commonly consists of three steps—the data is pre-processed, a measure of co-expression is calculated for every pair of genes, and a score cut-off is applied. Different approaches to data pre-processing and normalisation exist (Park *et al.*, 2003; Abbas-Aghababazadeh *et al.*, 2018). Further, after normalisation co-expression can be calculated in a number of different ways—for example, via a correlation coefficient or mutual information. A score cut-off is then usually imposed in order to identify gene pairs which are highly co-expressed. Alternatively, weighted networks can be analysed, in which edge weights correspond to levels of coexpression (Langfelder and Horvath, 2008).

There are many available methods for network construction which can be applied to the same data set, and which then lead to different networks. Enrichment analysis or comparison to orthogonal data, such as protein interaction data, is commonly used for network selection and validation. However, for many species and datasets poor or non-existent functional annotation makes this type of validation difficult. It is therefore often not clear which of the available network construction methods should be prioritised (De Smet and Marchal, 2010).

Here we introduce COGENT (COnsistency of Gene Expression NeTworks), an R package designed to aid the choice of a network construction pipeline without the need for annotation or external data. COGENT can be used to choose between competing co-expression measures, as well as to inform score cut-off choice. While designed for gene expression data, COGENT can be applied to other cases where network construction relies on similarity profiling, for example microbiome or synthetic lethality data.

## 2 Software description

### 2.1 Protocol schematic

Consistent co-occurrence of two gene products in the cell points towards a functional relationship between the genes. Gene product abundance, as well as co-occurrence, is a continuous-time phenomenon, which is experimentally observed at discrete time points or samples. The construction of a gene co-expression network can therefore be thought of as an estimation problem—we aim to infer general co-expression patterns from a limited set of data points. One way of investigating the success of such a procedure is through resampling. Networks constructed from a subset of all available samples will be noisier than the network constructed from the full dataset. However, they should still resemble each other: if subsetting the data results in networks with little to no overlap, then the network construction procedure may be too sensitive to noise in the data.

COGENT evaluates network construction methods through iterative resampling. At each step, the gene expression samples are split into two possibly overlapping sets of equal size. The same network construction function *f* (·) is applied to both sets in order to obtain two gene co-expression networks *G*_1_ = (*V, E*_1_) and *G*_2_ = (*V, E*_2_). COGENT then calculates several measures of consistency between these two networks.

The entire procedure is repeated multiple times in order to obtain robust results. Network construction methods which result in highly similar pairs of networks *G*_1_ ≈ *G*_2_ are considered to be consistent. When two or more competing methods are considered, the method exhibiting higher internal consistency should be preferred.

### 2.2 Edge set consistency

Consistency in COGENT is measured through a network comparison step at each iteration. The two networks *G*_1_ and *G*_2_ are considered to be similar if their edge sets are similar (*E*_1_ ≈ *E*_2_). We measure agreement between *E*_1_ and *E*_2_ using a (weighted) Jaccard index to produce two measures of edge set consistency—global and local similarity (Kao and Porter, 2018).

Since global similarity scales with network density, a density adjustment is required when methods resulting in different network densities are compared. This is particularly important when choosing a score cut-off. Density adjustment in COGENT is carried out through comparison to random networks generated using a configuration model.

### 2.3 Node metric consistency

Density-independent network comparison can also be performed by calculating a node metric such as degree or betweenness for all nodes in each of the two networks, and then comparing the obtained metric values. If the aim of downstream network analysis is, for example, to identify genes with many co-expression partners, the degree is a natural node metric to use.

At each COGENT iteration, a node metric set by the user can be applied to *G*_1_ and *G*_2_, resulting in two node metric vectors *d*_1_ and *d*_2_, respectively. The two can be compared in three different ways: via a correlation coefficient, rank k-similarity (Trajanovski *et al.*, 2013; Bozhilova *et al.*, 2019), and Euclidean distance.

## 3 Application

Global and local similarity and adjusted edge consistency, as well as node metric comparisons can all be used to evaluate network consistency. By iteratively resampling the data and measuring network consistency, COGENT can be used to prioritise different network construction pipelines, as well as to inform co-expression cut-offs without the need for external validation data. In Figure 1 we illustrate how COGENT can be used to select between Pearson and Kendall correlation coefficients for measuring co-expression, as well as how to select the score cut-off. A full worked example can be found in the SI. COGENT has also been used to assess signed distance correlation as a measure of gene co-expression (Pardo-Diaz *et al.*, 2020).

**Figure 1:**
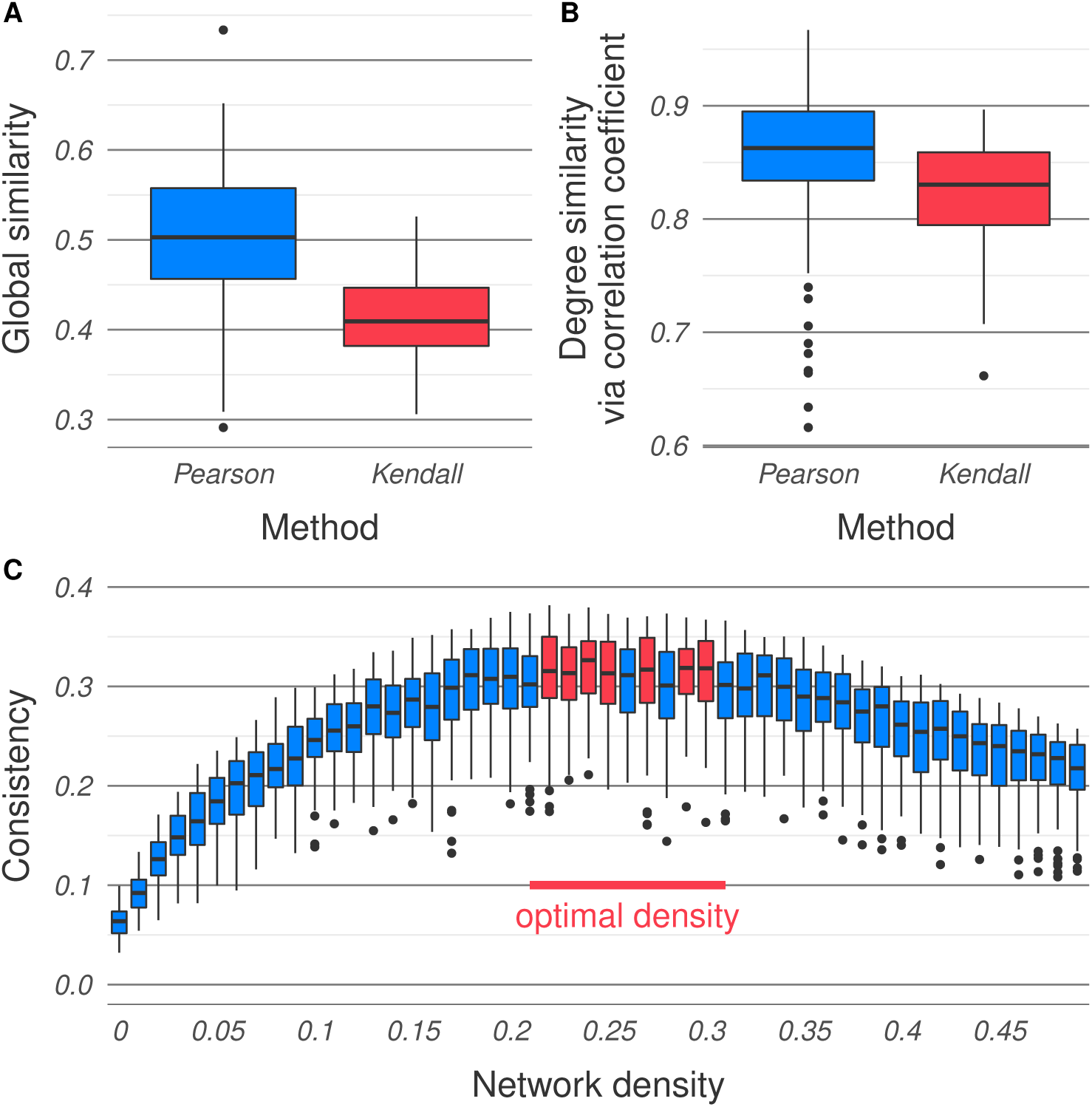
Sample output for data obtained from the Expression Atlas (accession number E-MTAB-5174, Petryszak *et al.*, 2015). See SI for details. (A) Global similarity indicates that Pearson correlation coefficients should be preferred over Kendall coefficients for measuring co-expression. (B) Consistency of the network degree sequences also indicates Pearson should be preferred. (C) The optimal edge density for Pearson networks is between 0.23 and 0.31.

While originally developed for gene expression data, COGENT can also be used for the inspection of other data types. For example, it can be applied to microbiome data in order to identify symbiotic organisms, or to synthetic lethality data in order to identify genetic interactions with high confidence.

## Supporting information

Software tutorial

