## Supplemental Information for "COGENT: evaluating the consistency of gene co-expression networks"

**Supplementary Information**

Lyuba V. Bozhilova[1], Javier Pardo-Diaz[1], Gesine Reinert[1], Charlotte M. Deane[1,*]
[1]Department of Statistics, University of Oxford, Oxford, OX1 3LB, United Kingdom
[*]

### 1 Introduction

The COGENT workflow is illustrated and explained in Section 2. Implementation details, as well as a complete list of functions available in COGENT are presented in Section 3. Network consistency measures used by COGENT are explained in detail in Section 4. Finally, an example application of COGENT is presented in Section 5. A similar, interactive application can be found in the tutorial.

### 2 Workflow

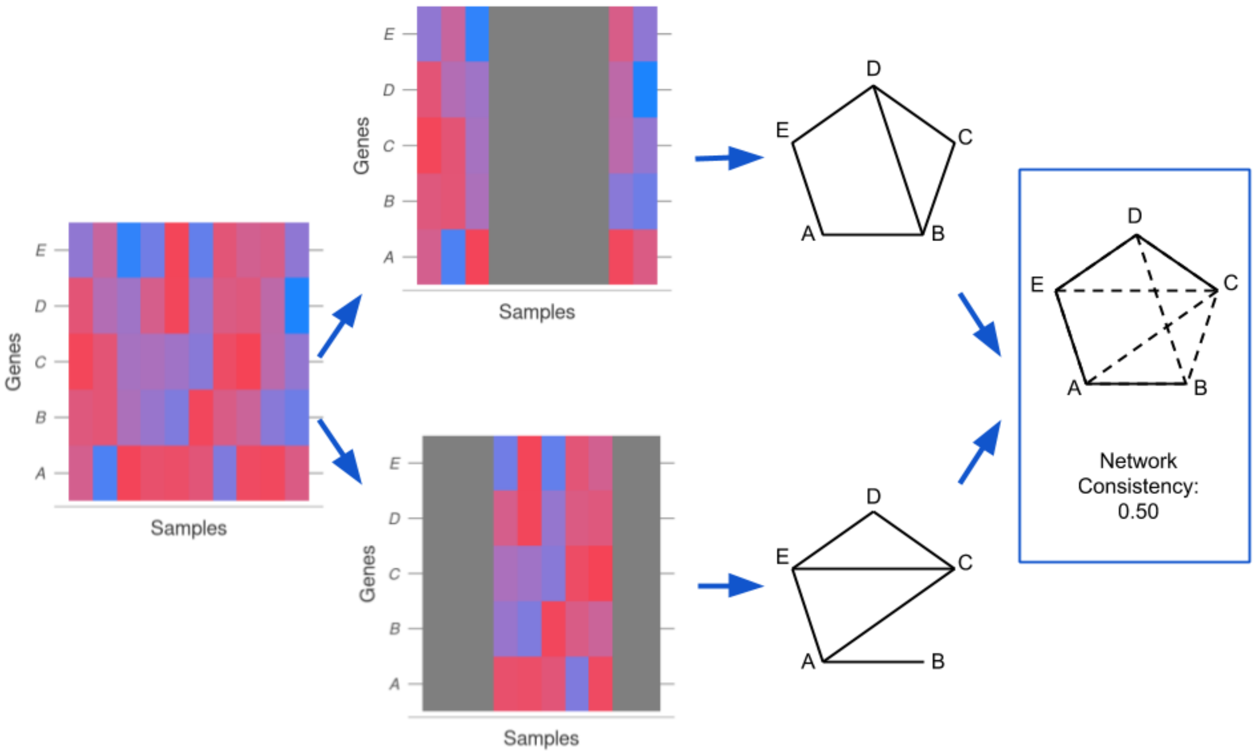


Figure 1: **COGENT workflow schematic.** In this example, the input data is a gene expression matrix with rows corresponding to genes—in this case $A, B, C, D, E$—and columns corresponding to samples (far left). First, the expression matrix columns are randomly split into two possibly overlapping groups of equal size (left). Then, a network is constructed from each of the sample groups (right). Finally, the two resulting networks are compared and the consistency between them is calculated (far right). In this example, the two networks have six edges each, and overlap at four of these edges. One measure of their consistency is the Jaccard index between their edge sets (see Section 4), which in this case is 0.50.

COGENT evaluates network construction methods through iterative resampling. At each step, the gene expression samples are split into two possibly overlapping sets of equal size. The overlap amount can be controlled by the user, and the sets are generated at random. In cases where a large number of samples are available, the subsets can be left disjoint.

After the two random gene expression subsets are generated, the same network construction method is applied to both in order to obtain two gene co-expression networks. Since the data is split by samples and not by genes, the two resulting networks should have identical or significantly overlapping node sets. COGENT then calculates several measures of consistency between these two networks. This workflow is illustrated in Figure 1.

The entire process is repeated multiple times in order to obtain robust results (see Algorithm 1). Consistency can be calculated in a number of different ways, chosen and parametrised by the user. Network construction methods which result in highly similar pairs of networks are considered to be consistent. When two or more competing methods are considered, the method exhibiting higher internal consistency should be preferred.

---

**Algorithm 1** COGENT workflow.

---

**Require:** Gene expression data $M$, network construction method $\psi(\cdot)$, number of repetitions $n$, [optional] node metric $f(\cdot)$
  **for** $i \in \{1, 2, \ldots, n\}$ **do**
    Randomly split $M$ into $M_1$, $M_2$
    Create networks $G_1 = \psi(M_1)$, $G_2 = \psi(M_2)$
    Calculate the edge set consistency $EC_i$ between $G_1$ and $G_2$
    **if** node metric $f(\cdot)$ is provided **then**
      Calculate the node metric consistency $NC_i$ between metric value vectors $\overrightarrow{f(V(G_1))}$ and $\overrightarrow{f(V(G_2))}$
    **end if**
  **end for**
  **return** $\overrightarrow{EC} = \{EC_1, \ldots, EC_n\}$ and [optional] $\overrightarrow{NC} = \{NC_1, \ldots, NC_n\}$

---

### 3 Software description

#### 3.1 Implementation

COGENT was built in R v.3.6.1 [11] using devtools [17] and roxygen2 [16]. It imports the core R package parallel (v.3.6.1 and higher), as well as the packages Matrix (v.1.2.17 and higher) and igraph (v.1.2.4.1 and higher), all of which are available on CRAN.

The main COGENT functions take as input gene expression data (an R object of class data frame) and a network construction method (an R object of class function) and return a set of consistency metrics. These consistency metrics aim to capture the suitability of the network construction method when applied to the data. Higher consistency implies a more robust network construction method.

#### 3.2 Function overview

The R package COGENT includes 15 functions, listed in Table 1. They are split into four categories—input checks, internal functions, network similarity (i.e. consistency) functions, and main COGENT functions.

#### 3.3 Input checks

The input check functions are `checkArrayList`, `checkExpressionDF`, `checkFun` and `checkMatrixList`. They are used to check whether the input (and output) of intermediate steps of the COGENT pipeline adhere to the required format.

Different types of common errors are clearly documented, so that input checks can be used for debugging before any computationally demanding analysis is performed.

#### 3.4 Internal functions

The internal functions include alignment functions (`alignArrays` and `alignMatrices`), similarity and distance functions (`calculateEuclideanDistance` and `calculateKSimilarity`) and the random data split function (`splitExpressionData`).

Network comparison in COGENT relies on a one-to-one mapping of nodes in the pair of networks generated at each iteration. If the network construction method does not respect the order in which

| Function | Description |
|---|---|
| alignArrays | Align node metric arrays |
| alignMatrices | Align adjacency matrices |
| calculateEuclideanDistance | Euclidean distance |
| calculateKSimilarity | Rank k-similarity |
| checkArrayList | Check node metric list |
| checkExpressionDF | Check a gene expression data frame |
| checkFun | Check function |
| checkMatrixList | Check adjacency matrix list |
| cogentLinear | Multiple COGENT calls, executed on a single thread |
| cogentParallel | Multiple COGENT calls, executed in parallel |
| cogentSingle | Single COGENT call |
| getEdgeSimilarity | Get the edge similarity for two networks |
| getEdgeSimilarityCorrected | Get the edge similarity with configuration model correction |
| getNodeSimilarity | Get the node similarity of two networks |
| splitExpressionData | Randomly subset expression samples |

Table 1: **Functions implemented in COGENT.**

genes are labelled in the expression data set, e.g. by removing isolated nodes, an additional mapping step is needed. Similarly, if the node metric function does not respect a fixed node ordering, additional mapping may be required before the node metric comparison step. This mapping, or alignment, is not performed by default, and appropriate error messages are produced if a mismatch is detected.

The distance and similarity functions are, respectively, Euclidean distance with optional rescaling to $[0, 1]$ and parametrised rank $k$-similarity [13]. For rank $k$-similarity, the top proportion of ranks to be compared can be given as a quantile (e.g. the top 10% of nodes) or as an integer (e.g. the top 100 nodes). See Section 4.3 for details.

#### 3.5 Network similarity

Network similarity functions include the edge set consistency functions calculated by `getEdgeSimilarity`, the density adjusted edge set consistency in `getEdgeSimilarityCorrected`, and node metric consistency functions calculated by `getNodeSimilarity`. Note that global and local similarity are calculated simultaneously by `getEdgeSimilarity`. Density adjustment in `getEdgeSimilarityCorrected` can be carried either with the fully random correction term, or with semi-random correction. Similarly, `getNodeSimilarity` calculates node metric consistency in any of the three available ways (via a correlation, rank $k$-similarity, or Euclidean distance). Details on the calculation of all of these are provided in Section 4.

#### 3.6 Main functions

The main COGENT functions `cogentSingle`, `cogentLinear` and `cogentParallel` execute the complete COGENT pipeline. The principal input of these are a gene expression data set, stored in a data frame, and a network construction method, which transforms the data frame to a network adjacency matrix.

The function `cogentSingle` carries out a single iteration of COGENT. It splits the data in two random subset, constructs a network from each dataset, and returns a number of consistency metrics (e.g. global similarity, average local similarity, and rank $k$-similarity for some user-defined node metric). Optional parameters, e.g. whether to do any alignment or what proportion of the data is shared during the data split step, are passed to the internal and network similarity functions described in Sections 3.4 and 3.5.

The functions `cogentLinear` and `cogentParallel` are wrappers for `cogentSingle` and are used for carrying out multiple COGENT iterations. The former executes COGENT linearly, and is suitable for cases where network construction is fast or slow, but parallelised. The latter executes COGENT in parallel on multiple threads. It is designed for cases where network construction is slow and not parallelised.

### 4 Network consistency

Consistency in COGENT is measured through a network comparison step at each iteration. Suppose we are interested in studying some gene expression data set $M_0$, and wish to construct a co-expression network $G_0$ from it by applying some function $G_0 = \psi(M_0)$ to the data. For example, $\psi(\cdot)$ may involve first calculating Pearson correlation coefficients between all pairs of gene expression profiles, i.e. all pairs of rows of $M_0$, and then selecting the top 10% of these as edges. This will result in a co-expression network, where the 10% most strongly co-expressed pairs of genes are linked by edges.

The first step of COGENT is to randomly split the data samples $M_0$ in two groups of equal size, $M_1$ and $M_2$. Note that the split is performed only over the samples, so both groups will contain data for all genes in $M_0$. By default, $M_1$ and $M_2$ do not overlap. However, a level of overlap can be set by the user if the overall number of samples in $M_0$ is low. The network construction function $\psi(\cdot)$ is then applied to $M_1$ and $M_2$ in order to produce two networks, $G_1 = \psi(M_1)$ and $G_2 = \psi(M_2)$. Consistency is measured by comparing $G_1$ and $G_2$. We first use edge overlap as a measure of consistency. We calculate a global edge overlap across the networks, and local overlap, which measures how well each gene neighbourhood is preserved across $G_1$ and $G_2$ [7]. Both of these can be applied to unweighted, as well as weighted networks. Since edge overlap is expected to be higher for denser networks, we introduce a density adjustment for the global edge overlap measure. The density adjusted consistency measure is only suitable for unweighted networks and can be used to inform threshold choice. Finally, we introduce an optional measure of node metric consistency. If, for example, genes of high betweenness in the co-expression network are of interest, then $G_1$ and $G_2$ can be compared by the betweenness centrality of their nodes, rather than by the overlap of their edges.

#### 4.1 Edge set consistency

Suppose that a single iteration of COGENT applied to some some data $M_0$ and network construction method $\psi(\cdot)$ results in two networks $G_1 = (V, E_1)$ and $G_2 = (V, E_2)$ with a shared node set $V$ and edge sets $E_1$ and $E_2$. The two networks $G_1$ and $G_2$ can be considered to be similar if $E_1 \approx E_2$. If such similarity is replicated across COGENT iterations, $\psi(\cdot)$ is a consistent method for constructing networks from the data $M_0$. We measure agreement between $E_1$ and $E_2$ using a Jaccard index if the networks are binary (i.e. unweighted), and a weighted Jaccard index if they are weighted.

##### 4.1.1 Unweighted and weighted Jaccard index

Let $A$ and $B$ be two finite non-empty sets. These could be, for example, the edge sets of two unweighted networks. The *Jaccard index* [6] between them is calculated as

$$Jacc(A, B) = \frac{|A \cap B|}{|A \cup B|}. \tag{1}$$

This is a generic metric of set overlap, and is not restricted to network comparison; it is also known as Tanimoto similarity [12, 2]. It is always in the range $[0, 1]$, with zero corresponding to no set overlap $A \cap B = \emptyset$ and one corresponding to perfect overlap $A \equiv B$.

Applied over edge sets, the Jaccard index can be used to provide a measure of both global and local network similarity, for example for community detection in multilayer networks [15, 7]. In COGENT, we generalise these to the weighted case and use them to perform network comparison between $G_1$ and $G_2$ at every iteration of our algorithm.

The Jaccard index can be used to evaluate the similarity of two edge sets, but does not take into account edge weights. A measure of set overlap which penalises weight difference is the *weighted Jaccard index*. Suppose that for some finite index set $\mathcal{I}$, we have two non-negative weight vectors $A = \{a_e\}_{e\in\mathcal{I}}$ and $B = \{b_e\}_{e\in\mathcal{I}}$. In the context of weighted networks, $a_e$ could be the edge weight of $e \in V^2$ in the network $G_1$ and $b_e$ could be the weight of $e$ in $G_2$. Non-edges are assumed to have zero weight. The weighted Jaccard index between the weight vectors $A$ and $B$ is

$$Jacc(A, B) = \frac{\sum_{e \in \mathcal{I}} min(a_e, b_e)}{\sum_{e \in \mathcal{I}} max(a_e, b_e)}. \tag{2}$$

Like the Jaccard index (Equation 1), the weighted Jaccard index is in the range $[0, 1]$, with higher values corresponding to better agreement between $A$ and $B$. Further, there is a correspondence between the two indices. If the weights are binary, then the weighted Jaccard index $Jacc(A, B)$ reduces to the unweighted $Jacc(A^*, B^*)$ between $A^* = \{e \in \mathcal{I} : a_e = 1\}$ and $B^* = \{e \in \mathcal{I} : b_e = 1\}$.

#### 4.1.2 Global similarity

Suppose $G_1 = (V, E_1)$ and $G_2 = (V, E_2)$ are binary, i.e. unweighted, and non-empty. The *global similarity* between them is defined as the Jaccard index between their edge sets:

$$\textit{global similarity}\ (G_1, G_2) = \frac{|E_1 \cap E_2|}{|E_1 \cup E_2|}. \tag{3}$$

It can be interpreted as the ratio between the number of edges of the intersection of $G_1$ and $G_2$ and the number of edges of their union. If the two networks are similar, then their would not be much larger than their intersection, resulting in a global similarity close to one.

This measure does not take into account edge weights. It would be unsuitable for comparing dense, possibly complete co-expression networks, in which non-negative edge weights correspond to gene co-expression. Suppose $G_1$ and $G_2$ are weighted, and that for $e \in V^2$, $w_1(e)$ is the weight of $e$ in $G_1$, and $w_2$ is the weight of $e$ in $G_2$. We treat non-edges as equivalent to zero-weight edges, i.e. $w_i(e) = 0 \iff e \notin E_i$.

Intuitively, the two weighted networks $G_1$ and $G_2$ are similar if corresponding edges have similar weights, i.e. if for most $e \in V^2$, $w_1(e) \approx w_2(e)$. To measure similarity between the networks, we use the weighted Jaccard index over the edge weights:

$$\textit{global similarity}\ (G_1, G_2) = \frac{\sum_{e \in V^2} \min(w_1(e), w_2(e))}{\sum_{e \in V^2} \max(w_1(e), w_2(e))}. \tag{4}$$

The unweighted global similarity is equivalent to the weighted one, where edges are all assigned equal weight one. More generally, in the unweighted case, a possible edge $e \in V^2$ contributes to the numerator of the fraction, i.e. the graph intersection, if it is present in both networks, and to the the denominator of the fraction, i.e. the graph union, if it is present in at least one of the networks. In the weighted case, the smaller weight contributes to the numerator—in both networks, the edge appears with weight at least $\min(w_1(e), w_2(e))$—and the larger weight contributes to the denominator—in at least one of the networks, a weight of $\max(w_1(e), w_2(e))$ is observed.

#### 4.1.3 Local similarity

Rather than considering the complete edge sets, we can calculate a *local similarity* specific to the neighbourhood of every node in the network. Such a measure is of interest when heterogeneous noise is present. This can be the case in gene expression data, where noise is sometimes assumed to have gene-specific variance [5]. If a gene $v$ is strongly affected by experimental noise, we would expect that its neighbourhood in a gene co-expression network to be more strongly dependent on the choice of samples used for network construction. In this case, in a single iteration of COGENT, the neighbourhoods $\Gamma_{(1)}(v)$ of $v$ in $G_1$ and $\Gamma_{(2)}(v)$ of $v$ in $G_2$ may differ more than the neighbourhoods of other genes. In the unweighted case [7], this is quantified by:

$$\textit{local similarity}\ (v; G_1, G_2) = \frac{|\Gamma_{(1)}(v) \cap \Gamma_{(2)}(v)|}{|\Gamma_{(1)}(v) \cup \Gamma_{(2)}(v)|}. \tag{5}$$

This is related to the global similarity (Equation 3), where instead of the complete edge sets, only the neighbourhoods of $v$ are considered. For the weighted case, the measure can be defined analogously to Equation 4.

$$\textit{local similarity } (v; G_1, G_2) = \frac{\sum_{u \in V, u \neq v} \min(w_1(uv), w_2(uv))}{\sum_{u \in V, u \neq v} \max(w_1(uv), w_2(uv))}. \tag{6}$$

Like global similarity, local similarity for a node is in $[0, 1]$, with higher values corresponding to better preserved networks. Local similarity can be used to identify particularly consistent—or inconsistent—parts of the network, and it can also be used to provide a global network summary by averaging over all nodes [7]. The average local similarity and the global similarity are related but not necessarily identical. Their relationship is similar to that between the average local clustering coefficient and the global clustering coefficient, in that the former treats all nodes equally, and the latter is more strongly influenced by high-degree nodes.

#### 4.1.4 Edge overlap and network density

Both global and local similarities are measured by scaling the edge overlap between two networks. However, edge overlap is affected by network density. Denser networks are more likely to share edges by chance than sparser ones. Consider, for example, two random graphs on 4 nodes, each of which has precisely one edge. Since there are $\binom{4}{2} = 6$ ways of placing the edge, the probability that the networks share an edge is $1/6$, and most of the time the graphs would have a global similarity of zero. If the two graphs instead had five randomly placed edges each, they would always share at least four edges, meaning their global similarity would be at least $2/3$. However, in both cases the overlap is entirely random—so in a sense neither pair should exhibit better "consistency".

Since COGENT works by repeatedly applying a network construction function $\psi(\cdot)$ to random subsets of the data $M$ of equal size, we would expect the pairs of graphs $(G_1, G_2)_j$ obtained across different iterations $j \in \{1, 2, \dots\}$ to have similar density. Hence, global and local similarity can be meaningfully compared across iterations. If we consider an alternative network construction function $\xi(\cdot)$ which produces networks $(H_1, H_2)$ of similar density to the ones produced by $\psi(\cdot)$, then we can also meaningfully compare $\psi(\cdot)$ to $\xi(\cdot)$. For example, if $\psi(\cdot)$ builds edges from the top 10% of highest Pearson correlations between gene expression profiles, and $\xi(\cdot)$ builds edges from the top 10% of highest Kendall correlations, then global similarity would be a good way of identifying which method is more consistent.

If, however, $\chi(\cdot)$ uses the top 20% of Pearson correlation coefficients instead, comparing $\psi(\cdot)$ and $\chi(\cdot)$ would be less straightforward. As $\chi(\cdot)$ produces denser networks, we would expect the global and local similarities at COGENT iterations for it to be higher. If $\chi(\cdot)$ outperformed $\psi(\cdot)$, it would not be clear whether this was due to the method being genuinely more consistent. Higher similarity scores for $\chi(\cdot)$ compared to $\psi(\cdot)$ may just arise from the increase in network density. In order to account for this, we introduce a density adjusted edge set consistency metric.

### 4.2 Density adjusted edge set consistency

Density adjustment is required in order to compare between network construction methods $\psi(\cdot)$ and $\chi(\cdot)$ which result in networks of considerably different edge densities. This is particularly relevant when COGENT is used to determine a threshold or score cut-off in the construction of binary networks—e.g. whether to take the top 10% or top 20% of most highly co-expressed gene pairs to build a network from. The density adjustment implemented in COGENT relies on a comparison to randomly generated networks. First we explain the fully random density adjustment and study its dependency on network density in fully randomised networks. We then introduce a semi-random adjustment, and show that both adjustments produce similar results when applied to synthetic data. Both procedures are available within COGENT, but we note that the semi-random procedure is more computationally efficient and should generally be preferred. Density adjustment is designed specifically for unweighted networks. In the case of weighted networks, a density adjustment can be carried out by transforming network weights to quantiles, and calculating weighted global and local similarities on the transformed values.

#### 4.2.1 Fully random density adjustment

Consider a single COGENT iteration, resulting in two binary networks $G_1 = (V, E_1)$ and $G_2 = (V, E_2)$. Let $d_1$ and $d_2$ be the degree sequences of $G_1$ and $G_2$ respectively, so $d_1(v)$ is the degree of $v$ in $G_1$, and $d_2(v)$ is the degree of the same node $v$ in $G_2$.

Intuitively, we would like to know how much of the overlap between $E_1$ and $E_2$ is "genuine", and how much of it we might expect from any two networks of similar degree sequences. Density adjustment in COGENT corrects for the effect of random overlap by generating two networks, $G_1^* = (V, R_1)$ and $G_2^* = (V, R_2)$ from a type of configuration model, with random permutations $d_1^*$ and $d_2^*$ of the degree sequences $d_1$ and $d_2$ respectively. The degree sequences are permuted, since in the original network nodes correspond to genes and are therefore labelled. The networks $G_1^*$ and $G_2^*$ are generated with fixed degree sequences, no loops and no multiple edges, using the implementation by Csardi and Nepusz [4]. We define a correction term based on two pairwise comparisons between the co-expression and randomly generated networks: one between $G_1$ and $G_2^*$, and one between $G_2$ and $G_1^*$:

$$\alpha(G_1, G_2|G_1^*, G_2^*) = \frac{1}{2}(|E_1 \cap R_2| + |E_2 \cap R_1|). \tag{7}$$

Heuristically, the value of $\alpha$ corresponds to a level of intersection between $G_1$ and $G_2$ we may expect at random. Each of the terms in the sum is the overlap between one "real" co-expression network and a random network which resembles its counterpart. To define the *adjusted consistency* between $G_1$ and $G_2$ we subtract $\alpha(G_1, G_2|G_1^*, G_2^*)$ from the graph intersection and add it to the graph union of $G_1$ and $G_2$ in the unweighted global similarity (Equation 3), to give what we call the *fully random adjusted consistency*:

$$\alpha\text{-}adjusted\ consistency\ (G_1, G_2|G_1^*, G_2^*) = \frac{|E_1 \cap E_2| - \alpha(G_1, G_2|G_1^*, G_2^*)}{|E_1 \cup E_2| + \alpha(G_1, G_2|G_1^*, G_2^*)}. \tag{8}$$

Unlike global and local similarity, the density adjusted consistency measure can be negative. The lowest possible adjusted consistency is obtained when $G_1$ and $G_2$ share no edges, but $G_1^* \equiv G_2$ and $G_2^* \equiv G_1$. In this case, the adjusted consistency is $-1/3$. Perfectly overlapping $G_1$ and $G_2$ will have adjusted consistency close to 1, with higher consistency for lower density networks. If network overlap is random, then $|E_1 \cap E_2| \approx \alpha(G_1, G_2|G_1^*, G_2^*)$ and the adjusted consistency would be close to 0.

We illustrate that the fully random consistency in Equation 8 does not increase with density like the global and local similarities with a simulation study. We generated pairs networks $G_1$ and $G_2$ from the $\mathcal{G}(N, p)$ model on $N = 100$ nodes and with edge probability $p \in \{0.01, 0.02, \ldots, 0.50\}$. For each value of $p$ we generated 100 pairs $(G_1, G_2)$ and calculated the global similarity and the fully random adjusted consistency of each one. This resulted in a total of 5000 pairs of networks. Since the networks are random, we observed adjusted consistencies close to zero at all values of $p$. Further, there was no relationship between the adjusted consistencies and $p$ (Pearson correlation coefficient 0.025). In contrast, the global similarity increased with $p$ (Pearson correlation coefficient 0.994). See Figure 2 for details.

Generating the pair of random networks $G_1^*$ and $G_2^*$ can be computationally expensive when the number of nodes in the network is large. This is due to the random graph generation algorithm: first, "stubs" are attached to every node in the network, and then pairs of stubs are randomly connected to build edges. However, every time the procedure creates a self-edge (two stubs of the same node are connected), or a multi edge (two stubs are connected between the pair of nodes which already share an edge), the process is restarted. In order to speed up the comparison, we introduce a semi-random density adjustment, based on approximate expectations of edge occurrence rather than on randomly generated networks.

#### 4.2.2 Semi-random density adjustment

In Equation 7 we introduced a density adjustment term by randomising the degree sequences of $G_1 = (V, E_1)$ and $G_2 = (V, E_2)$ and creating two random networks $G_1^* = (V, R_1)$ and $G_1^* = (V, R_2)$ with the same degree distributions as $G_1$ and $G_2$ respectively. We then calculated the graph intersection

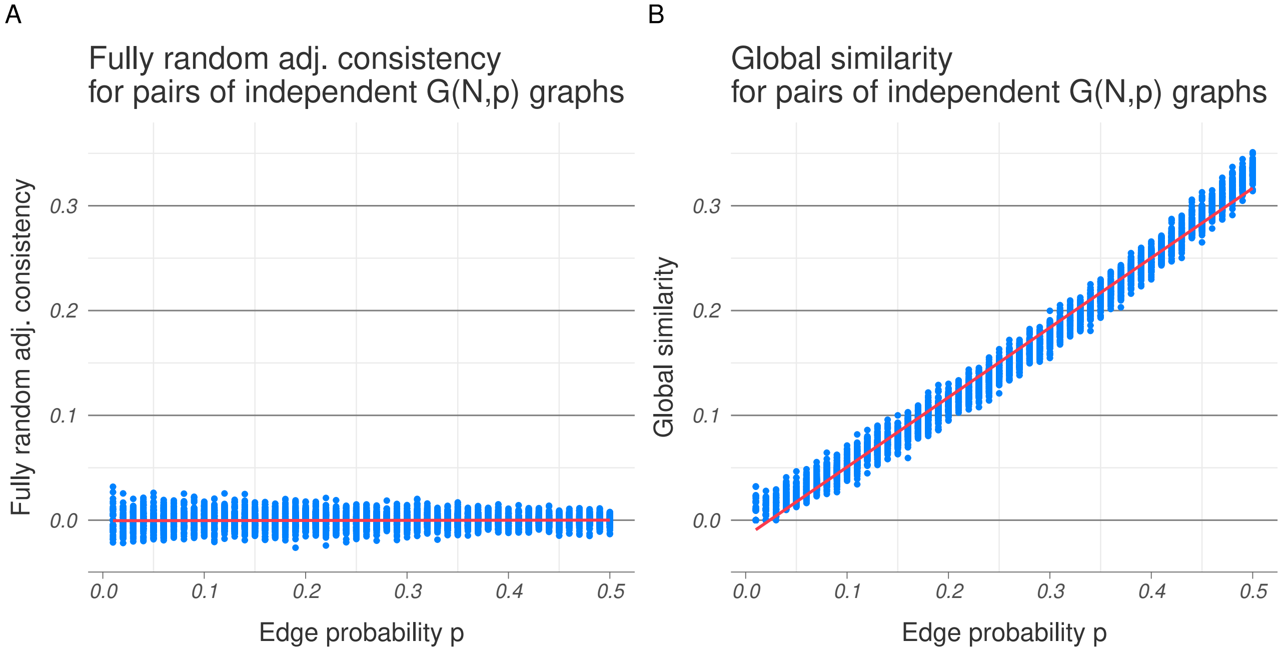


Figure 2: **Fully random adjusted consistency of $\mathcal{G}(N,p)$ pairs.** For different values of $p$, one hundred pairs of networks were generated from $\mathcal{G}(100,p)$ and the fully random adjusted consistency (A) and global similarity (B) were calculated for each pair. The dots (blue) correspond to the obtained values and the lines (red) were fitted using least squares regression. Consistency remained close to zero far all network densities, while global similarity increased with $p$.

between the fixed $G_1$ and the random $G_2^*$, as well as between $G_2$ and $G_1^*$. These intersections are both also random graphs by construction. Based on the intersections, we calculated a correction term $\alpha$.

Rather than generating the random graph intersections, we can calculate a correction term $\beta$ analogous to $\alpha$ by approximating their expected size. Without loss of generality, consider the intersection between $G_1$ and $G_2^*$. First, note that if an edge $e = (u,v)$ is not present in $G_1$, i.e. $e \notin E_1$, then it clearly cannot be present in the intersection either. Therefore, with $G_1$ fixed,

$$\mathbb{P}(e \in E_1 \cap R_2) = \mathbb{1}\{e \in E_1\}\mathbb{P}(e \in R_2). \tag{9}$$

The graph $G_2^*$ is generated from a configuration model, after discarding any networks with self-loops and multi-edges. However, for sparse, large networks, the density of self-loops and multi-edges in the configuration model is low. Therefore, we can approximate the probability of an edge $e = (u,v)$ existing in $G_2^*$ with the probability of edge occurrence in the standard Molloy–Reed configuration model, which is proportional to $d_2^*(u)d_2^*(v)$ [9]. We normalise so that the expected number of edges in the network is $|R_2| = |E_2| = \frac{1}{2}\sum_u d_2^*(u)$ to get

$$\mathbb{P}(e = (u,v) \in R_2 | d_1^*, d_2^*) \approx \frac{d^*(u)d^*(v)}{K} \textit{ for } u \neq v, \tag{10}$$

where the normalising constant $K$ is

$$K = \frac{1}{|R_2|}\sum_{u>v} d^*(u)d^*(v) = \frac{1}{|E_2|}\sum_{u>v} d(u)d(v). \tag{11}$$

Given a random degree permutation $\{d_2^*(u)\}_{u\in V}$, the probability in Equation 9 for $u \neq v$ can be approximated by

$$\mathbb{P}(e = (u,v) \in E_1 \cap R_2 | d_1^*, d_2^*) \approx \mathbb{1}\{e \in E_1\}\frac{|E_2|d_2^*(u)d_2^*(v)}{\sum_{u>v} d_2^*(u)d_2^*(v)} =: p_e. \tag{12}$$

This allows us to calculate an approximation to the expected adjacency matrix of the intersection between $G_1$ and $G_2^*$, given the degree permutation used for generating $G_2^*$. This adjacency matrix can be thought of as a weighted network, where edge weights $p_{uv}$ correspond to probability of edge occurrence. The expected number of edges in the overlap is then approximately the sum of all $p_{uv}$:

$$O_{12} = \sum_{e} p_e \propto \sum_{e=(u,v)\in E_1} d_2^*(u)d_2^*(v). \tag{13}$$

We can approximate the overlap $O_{21}$ between $G_2$ and $G_1^*$ analogously, replacing the degree sequence $d_2$ with a random permutation $d_1$ of the degree sequence of $G_1$. Using both, we define a new correction term $\beta$:

$$\beta(G_1, G_2|d_1^*, d_2^*) = \frac{1}{2}(O_{12} + O_{21}). \tag{14}$$

The new correction term $\beta(G_1, G_2|d_1^*, d_2^*)$ approximates the expectation of $\alpha(G_1, G_2|G_1^*, G_2^*)$ given that the degree sequence permutations $d_1^*$ and $d_2^*$ were used to generate $G_1^*$ and $G_2^*$ respectively. We call this correction term "semi-random" because while it does not require the generation of the graphs $G_1^*$ and $G_2^*$, it still requires the random permuting of the degree sequences $d_1$ and $d_2$. Finally, we use $\beta$ to define a *semi-random adjusted consistency* measure, similar to the fully random one in Equation 8:

$$\beta\text{-}adjusted\ consistency\ (G_1, G_2|d_1^*, d_2^*) = \frac{|E_1 \cap E_2| - \beta(G_1, G_2|d_1^*, d_2^*)}{|E_1 \cup E_2| + \beta(G_1, G_2|d_1^*, d_2^*)}. \tag{15}$$

In its implementation, the semi-random adjusted consistency relies entirely on matrix algebra and array permutations, rather than on network generation. This makes it more computationally efficient than the fully random one.

To evaluate its performance, semi-random adjusted consistency was measured for the set of $\mathcal{G}(N, p)$ pairs of networks discussed in the previous section. Like the fully random adjusted consistency, its values were close to zero and did not increase with network density. Further, they correlated well with the fully random values (Pearson correlation coefficient 0.807). Therefore, the approximations made are appropriate. See Figure 3 for details.

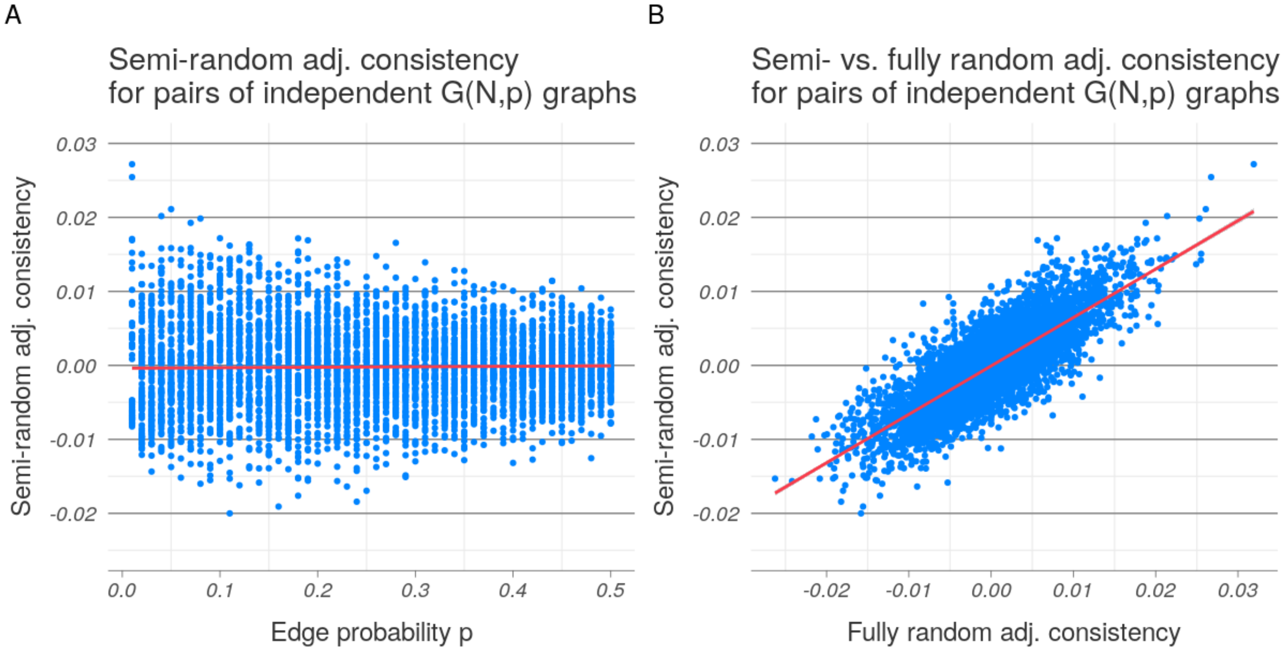


Figure 3: **Semi-random adjusted consistency of $\mathcal{G}(N, p)$ pairs.** (A) Like the fully random adjusted consistency (Figure 2A), the semi-random measure remained close to zero as the edge probability $p$ increased. (B) The fully and semi-random measures were well-correlated. The dots (blue) correspond to the obtained values and the lines (red) were fitted using least squares regression.

Overall, we have demonstrated that both the fully random and the semi-random density adjusted consistency measures can be used to measure the agreement between two binary networks, controlling for their densities. This makes them suitable for comparing network construction algorithms resulting in binary networks of significantly different densities. Like global and local similarity, density adjusted consistency is a measure of edge overlap. Within COGENT, we introduce a further way of evaluating network consistency using node metric agreement.

#### 4.3 Node metric consistency

Edge overlap is not the only way to think about network similarity. For example, consider two transport networks of Britain, where nodes correspond to villages, towns and cities, and edges are either roads or direct railway connections. We may argue that the railway network is in a sense functionally similar to the road network, since both identify the same large cities as major transport hubs. This similarity is between the node properties of the two networks, and is not based on edge overlap. It also depends on the node metric of choice—while the high-degree nodes might be the same across the two networks, the high betweenness nodes may be different.

Both degree and betweenness centrality as well as other node metrics are used in co-expression network analysis to identify key functionally relevant genes, e.g. [14]. It therefore may be desirable to study the consistency of a network construction method with respect to these metrics. That is, we may ask how much we can trust node metric values obtained from the co-expression network, rather than how reliable the network itself is. Node metric consistency can be calculated in three different ways within COGENT.

Suppose again that a COGENT iteration results in a pair of networks $G_1 = (V, E_1)$ and $G_2 = (V, E_2)$, and let $f : V(G) \to \mathbb{R}$ be a node metric of interest. A network construction method may be considered consistent *with respect to the node metric* $f$ if $f(v|G_1) \approx f(v|G_2)$ for all $v \in V$. COGENT can measure the agreement between $f(v|G_2)$ and $f(v|G_2)$ in three different ways:

- through Euclidean distance,
- through a correlation coefficient, and
- through rank $k$-similarity [13, 3].

The Euclidean distance measures the distance between the vectors $f(V(G_1))$ and $f(V(G_2))$ in $\mathbb{R}^V$ space. The obtained value is non-negative, with *lower* values corresponding to better network consistency. Optionally, a scaled Euclidean distance can be calculated, for which metric values are first scaled to the range $[0, 1]$ using

$$f^*(u) = \frac{f(u) - \min_{v \in V}\{f(v)\}}{\max_{v \in V}\{f(v)\} - \min_{v \in V}\{f(v)\}} \tag{16}$$

and then the Euclidean distance between $f(V(G_1))^*$ and $f(V(G_2))^*$ is itself scaled between 0 and 1 using

$$d(f^*(V(G_1)), f^*(V(G_2))) = \frac{||f^*(V(G_1)) - f^*(V(G_1))||}{\sqrt{|V|}}. \tag{17}$$

This results in a measure between 0 and 1, where again *lower* values correspond to higher network consistency.

The correlation between the node metric vectors $f(V(G_1))$ and $f(V(G_2))$ can also be calculated. By default, Pearson correlation coefficients are calculated, and missing data is handled by pairwise deletions. That is, only nodes where a metric value was obtained in both $G_1$ and $G_2$ are used. However, in our implementation options are directly passed to the R-base correlation function $cor(\cdot)$, which allows Pearson, Spearman and Kendall correlation coefficients to be calculated, and missing data to be handled in different ways.

Rank $k$-similarity [13] can also be used to evaluate network consistency within COGENT. Briefly, rank $k$-similarity measures how well the sets of highest ranking nodes with respect to the metric $f(\cdot)$ in $G_1$ and $G_2$ agree. For example, it can be used to say what proportion of the top 50 highest degree nodes in $G_1$ are also among the highest degree nodes in $G_2$. The default value of $k$ in COGENT is 10% of the number of genes $|V|$.

Note that, since $k$-similarity is rank-based, it is also not dependent on the network density. More generally, if the node metric $f(\cdot)$ used or the consistency measure applied to it is not dependent on network density, node metric consistency can be used instead of density adjusted edge set consistency to choose between network construction methods of different densities.

Together, the different edge set and node metric consistency measures provide a profile of the network similarity between $G_1$ and $G_2$ at every COGENT iteration. By aggregating across multiple iterations, COGENT outputs a consistency profile of a network construction method $\psi(\cdot)$ when applied to a gene expression data set $M$. In the next section we illustrate how COGENT can be used to inform network construction in different cases. We focus on unweighted networks; applications with weighted networks are analogous.

### 5 Example application

The network comparison measures described in Section 4 can be used to evaluate the consistency of a network construction method $\psi(\cdot)$ as applied to a gene expression data set $M$. Comparing the consistency of different methods $\psi(\cdot)$ and $\xi(\cdot)$ allows the user to prioritise a construction method for further analysis. This prioritisation can happen with respect to overall network consistency, or with respect to a particular application. For example, if genes of high betweenness centrality are of interest, network consistency can be measured with respect to betweenness centrality. Our method allows the user to assess network quality without the use of any additional data, such as protein interaction or functional annotation data.

In this section we illustrate how COGENT can be used in two different scenarios—to choose between measures of gene co-expression for network construction and to set a co-expression score cut-off. In both applications we focus on a gene expression dataset obtained from the Expression Atlas [10].

#### 5.1 Gene expression data

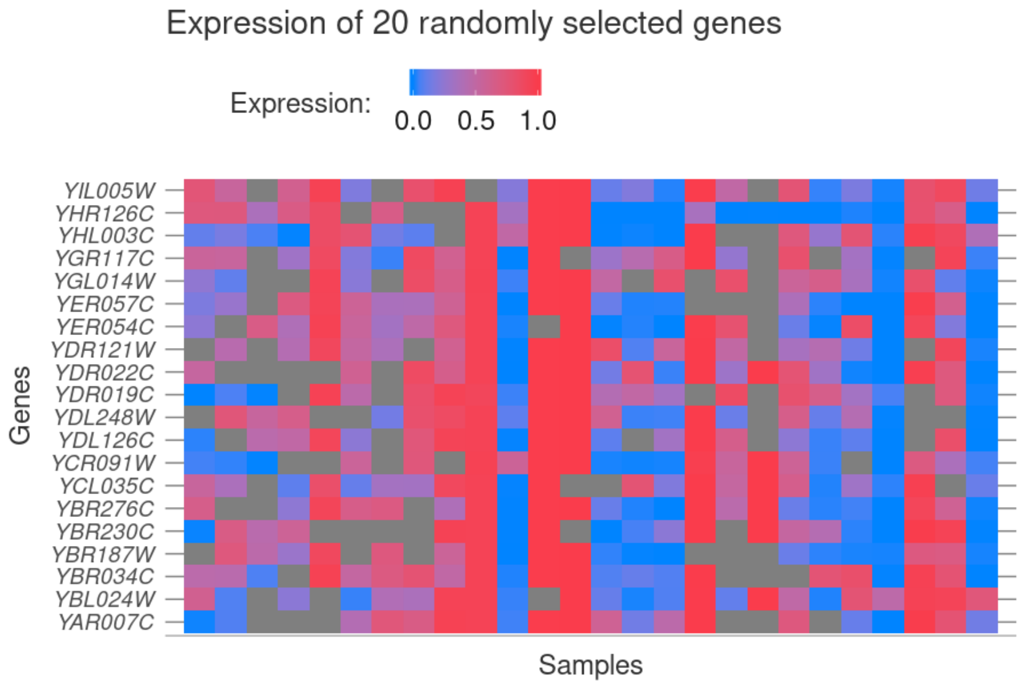


Figure 4: **Sample gene expression data for yeast.** Rows correspond to genes and columns correspond to samples. Expression values are scaled between 0 and 1, with low expression shown in blue and high expression shown in red. Missing values are shown in grey. Rows with similar colour patterns indicate high gene co-expression.

RNA-seq data of *Saccharomyces cerevisiae* (yeast) expressing pathways designed to increase ATP or GTP consumption was obtained from the Expression Atlas (accession number E-MTAB-5174). The original experiment as deposited in ArrayExpress [1] contains 156 samples, sequenced using Illumina HiSeq 2500. The curated data obtained from the Expression Atlas was reduced to expression profiles for 3103 genes across 26 samples. We further removed any genes where expression data was missing for over 25% of all samples, resulting in a dataset of 1920 gene expression profiles. Finally, we took the 500 genes with highest variation in gene expression as measured by the mean absolute deviation. This resulted in a final dataset $M$ of 500 gene expression profiles across 26 samples. Expression values for 20 randomly selected genes are shown in Figure 4.

#### 5.2 Choosing between measures of co-expression

We consider two simple ways of building gene co-expression networks from the data described in Section 5.1. The first builds a network by calculating Pearson correlation coefficients between expression profiles and creating edges between the gene pairs with highest co-expression. This is sometimes known as hard thresholding [8]. In the first instance, we take the top 5% of the co-expression values between pairs of genes, which results in a gene co-expression network on 500 nodes and 6238 edges. Note that 5% of all possible gene pairs is $5\%\binom{500}{2} = 6237.5$. A total of 49 nodes in the network are isolated.

The second construction method uses Kendall correlation coefficients instead of Pearson correlation coefficients. Again, we take the top 5% of obtained values to correspond to edges, resulting in a second network on 500 nodes and 6238 edges. Of the 58 isolated edges in the Kendall network, 43 are also isolated in the Pearson network.

We can use COGENT both to examine the consistency of a network construction method and to compare the Pearson and Kendall networks. The global similarity between the two networks is 0.57, corresponding to an overlap of 4512 edges, or 72% of the edges in each network. Edge differences are not spread uniformly across the networks—higher degree nodes generally have higher local similarity and nodes of extremely low degree in either network account for peaks at 0 and 1 (see Figure 5).

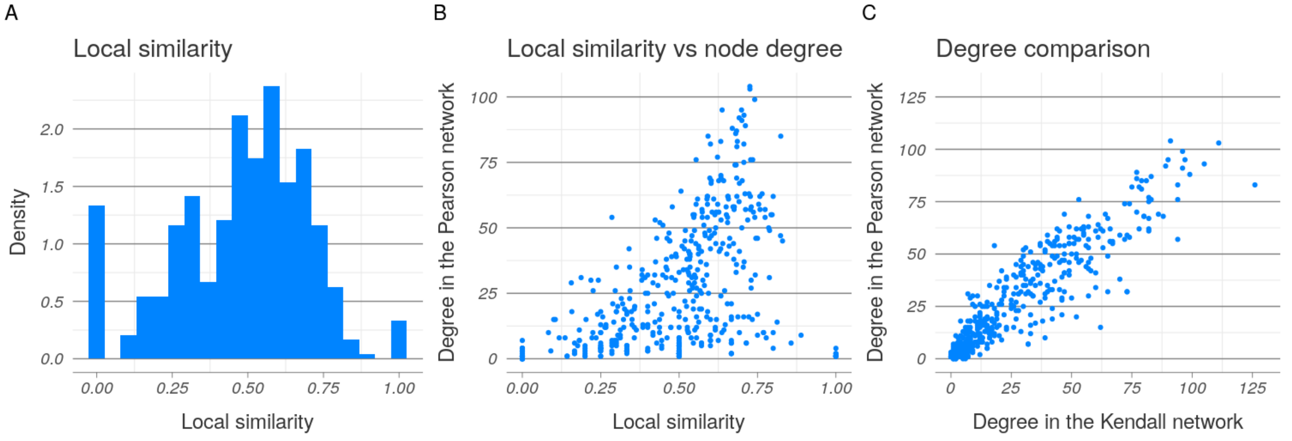


Figure 5: **Similarities between the Pearson and Kendall co-expression networks.** (A) Local similarity for each node (mean 0.48, st. dev. 0.22). Most values are symmetric around the mean, with additional peaks around 0 and 1. Binwidth is set to 0.05. (B) Local similarity plotted against node degree in the Pearson network. The peaks at 0 and 1 correspond to nodes of low degree. However, higher degree nodes tend to exhibit higher similarity. (C) The two degree distributions are positively correlated (Pearson correlation coefficient 0.92).

While the Pearson and Kendall networks share a significant number of edges, they are considerably different. We use COGENT to determine which of the two network correlation calculations is more consistent with respect to data resampling and which network should therefore be prioritised for further analysis. Since the networks are of the same density 0.05 by construction, density adjustment is not required. For illustration purposes, we evaluate edge set consistency through global similarity and node metric consistency for the degree through Pearson's correlation coefficient.

Since the dataset contains only 26 samples, we run COGENT for both methods with a sample overlap of 50% of the data at each iteration. Thus, when the data $M$ is resampled and split into subsets $M_1$ and $M_2$, 13 samples are randomly selected and included in both sets, and the remaining samples are randomly allocated to only one of the sets. One hundred COGENT iterations are used to evaluate the stability of each method. The computational complexity of the COGENT pipeline is linear in the complexity of the network construction method (and the node metric calculation). Thus, medium-to-large Pearson correlation networks can be analysed without any parallelisation on a standard desktop computer. COGENT has an additional inbuilt parallel mode, which is employed for the Kendall networks, where computing correlation matrices is slower.

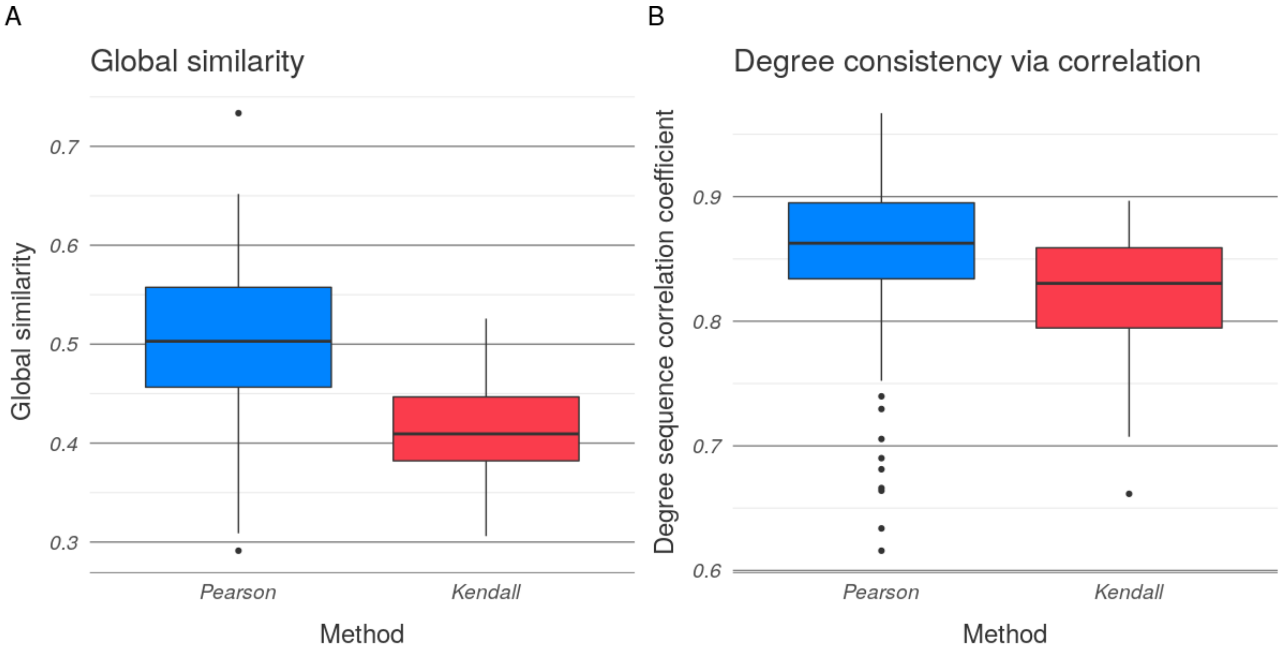


Figure 6: **COGENT analysis for Pearson and Kendall co-expression networks.** (A) Global similarity is visibly higher for the Pearson network construction method (mean 0.50, st. dev. 0.08) than for the Kendall method (mean 0.41, st. dev. 0.05). (B) While Pearson (mean 0.85, st. dev. 0.07) still outperforms Kendall (mean 0.82, st. dev. 0.04) in rank $k$-similarity for the degree, the difference is slightly less pronounced. This may be expected, since the two networks have similar degree distributions (Figures 5B). Note the difference in scales on the y-axis.

Edge set consistency as measured by global similarity is higher for the Pearson than the Kendall method (Figure 6). Pearson also outperforms Kendall in rank $k$-similarity for the degree with the default settings. While the difference is less pronounced for $k$-similarity than it is for global similarity, both are statistically significant. A two-sided Wilcoxon rank sum test for global similarity has p-value $p \approx 2.3 \times 10^{-15}$, and the same test for degree correlations has p-value $p \approx 4.2 \times 10^{-7}$.

The comparison between network consistency for both Pearson and Kendall correlation networks therefore indicates that the Pearson network should be preferred. However, this analysis was performed only on networks thresholded to ensure a 5% network density. In the next section, we use COGENT to study threshold choice.

#### 5.3 Imposing a co-expression score cut-off

In the previous section, we used COGENT to compare the consistency of two methods which both produced networks of 5% edge density. However, the value of 5% was arbitrary. We can also use COGENT in order to determine whether a different value may result in more consistent networks. We illustrate this by exploring different thresholds for a Pearson correlation network constructed using the same gene expression data.

As discussed in Figure 2, global similarity cannot be used to distinguish between methods with different network densities, because denser networks are more likely to have higher edge overlap by chance. In order to correct this, we use the semi-random density-adjusted edge set consistency measure in COGENT. We consider 50 density thresholds between 0.01 and 0.50, i.e. from taking only the top 1% of highest pairwise gene expression correlations down to taking the top half of these. At each threshold, we run the COGENT pipeline for 100 iterations, and split the data the same sample overlap as before (13 out of 26 shared samples in each sample subset).

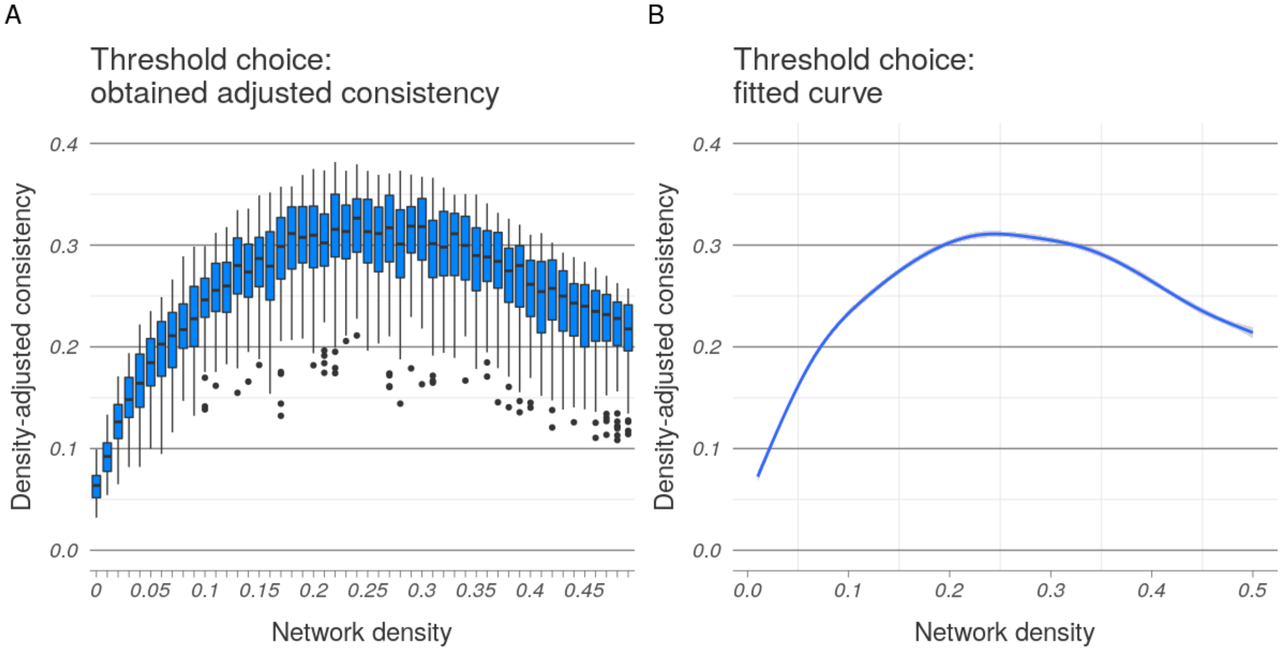


Figure 7: **COGENT analysis for Pearson thresholds.** (A) Obtained density adjusted edge set consistencies at different network densities. (B) Curve fitted to the data using a generalised additive model. The curve reaches its maximum around density 0.25, suggesting taking the top 25% highest correlation values may be optimal.

The density adjusted edge set consistency for the Pearson network at the 5% density threshold has mean 0.16 and standard deviation 0.03. As expected due to the semi-random correction term $\beta$, this is lower than the global similarity calculated using the same parameters (mean 0.50, standard deviation 0.08). As the network density increases, the adjusted similarity also increases until it plateaus for densities around 25%. For higher densities, the adjusted consistency then decreases (Figure 7). Fixing the network density around 25% may therefore be most preferable.

The highest observed adjusted consistency is at network density 25% (mean 0.32 and std. dev. 0.04). This is a significant improvement over the value at 5% density. However, differences in the plateau region are not necessarily statistically significant—a one-sided Wilcoxon's rank test ($\alpha = 5\%$) showed six other thresholds between 0.23 and 0.31 resulted in consistency values which were not significantly lower than those at the optimal threshold 0.25. Therefore, while 25% density results in networks of the highest consistency, COGENT can be used to identify a range of permissible thresholds.

We have illustrated the two main expected uses of COGENT with respect to gene co-expression data: to choose between different measures of co-expression, and to inform threshold choice for binary networks. Both of these problems arise when networks are constructed from other types of occurrence data.

[4] Gabor Csardi and Tamas Nepusz. The igraph software package for complex network research. *InterJournal*, Complex Systems:1695, 2006.

[5] Chuan Gao, Ian C McDowell, Shiwen Zhao, Christopher D Brown, and Barbara E Engelhardt. Context specific and differential gene co-expression networks via Bayesian biclustering. *PLoS Computational Biology*, 12(7):e1004791, 2016.

[6] Paul Jaccard. Nouvelles recherches sur la distribution florale. *Bulletin de la Societe Vaudoise des Sciences Naturelles*, 44:223–270, 1908.

[7] Ta-Chu Kao and Mason A Porter. Layer communities in multiplex networks. *Journal of Statistical Physics*, 173(3-4):1286–1302, 2018.

[8] Peter Langfelder and Steve Horvath. WGCNA: an R package for weighted correlation network analysis. *BMC Bioinformatics*, 9(1):559, 2008.

[9] Mark EJ Newman. *Networks*. Oxford University Press, Second edition, 2018.

[10] Irene Papatheodorou, Nuno A Fonseca, Maria Keays, Y Amy Tang, Elisabet Barrera, Wojciech Bazant, Melissa Burke, Anja Füllgrabe, Alfonso Muñoz-Pomer Fuentes, Nancy George, et al. Expression atlas: gene and protein expression across multiple studies and organisms. *Nucleic Acids Research*, 46(D1):D246–D251, 2017.

[11] R Core Team. *R: A Language and Environment for Statistical Computing*. R Foundation for Statistical Computing, Vienna, Austria, 2019.

[12] Taffee T Tanimoto. IBM internal report. *Nov*, 17:1957, 1957.

[13] Stojan Trajanovski, Javier Martín-Hernández, Wynand Winterbach, and Piet Van Mieghem. Robustness envelopes of networks. *Journal of Complex Networks*, 1(1):44–62, 2013.

[14] Sipko van Dam, Urmo Vosa, Adriaan van der Graaf, Lude Franke, and Joao Pedro de Magalhaes. Gene co-expression analysis for functional classification and gene–disease predictions. *Briefings in Bioinformatics*, 19(4):575–592, 2017.

[15] András Vörös and Tom AB Snijders. Cluster analysis of multiplex networks: Defining composite network measures. *Social Networks*, 49:93–112, 2017.

[16] Hadley Wickham, Peter Danenberg, and Manuel Eugster. *roxygen2: In-Line Documentation for R*, 2018. R package version 6.1.1.

[17] Hadley Wickham, Jim Hester, and Winston Chang. *devtools: Tools to Make Developing R Packages Easier*, 2019. R package version 2.2.1.
