## Supplementary material for "COGENT: evaluating the consistency of gene co-expression networks": Software tutorial

### COGENT tutorial

This is an R Markdown notebook. The .Rmd version of this file works like a Jupyter notebook – you can either click *Run* on each code chunk, or *Ctrl+Shift+Enter* your way through. You can find different versions of this tutorial (.Rmd, .pdf and .html) as part of the COGENT package on GitHub.

#### Introduction

If two genes have similar expression patterns, i.e. if their products appear in the cell at the same time, this could be evidence of shared biological function. Such information is often captured by gene co-expression networks. In these, genes are represented by nodes, and pairs of genes which are highly co-expressed are linked by edges.

There are a number of ways such networks can be built from gene expression data across multiple samples. One possibility is to calculate a Pearson correlation coefficient between the expression profiles of every pair of genes, and then build edges where the correlations are high (say, take the top 10% of values). Various choices of similarity metrics and thresholds are possible here, and there exist more complex network construction approaches. In the absence of external validation data, it is not always clear which to network construction method to apply. Therefore, a principled approach to method choice is needed.

In order to aid data-driven approaches to network construction, we develop the `R` package **COGENT** (**CO**nsistency of **G**ene **E**xpression **N**e**T**works). COGENT provides a suite of assessment tools for gene co-expression network construction. It can be used to choose between different network construction approaches, as well as to inform threshold choice. The main idea behind COGENT is that a good network construction function should produce similar networks when only a subset of the available expression samples are used.

#### Software requirements and installation

This is an `R` tutorial. The code and any associated files are available on GitHub as part of the COGENT package. The code can be run from terminal, through the RGui, or through RStudio. In order to execute the .Rmd notebook as is, you will need to use RStudio.

While the tutorial will work on any major operating system, we recommend using either Linux or macOS. Since the fork system call is not supported for Windows, you will not be able to execute parallelised code on Windows. You will have to run the code on a single thread instead.

Before we begin, make sure your `R` version and any already installed packages are up to date. COGENT was built using `R 3.6.1` but we recommend using the most recent version (`R 3.6.2` at the time of writing). In order to manage your pre-installed packages, run `old.packages()` to see a list of out-of-date ones, and run `update.packages()` to update them.

COGENT is available on GitHub, and the easiest way to install it is through `devtools::install_github`. You may need to install `devtools` first. Once you have devtools installed, you can uncomment and run the following to install COGENT:

```
# require("devtools")
# install_github("lbozhilova/COGENT")
```

This tutorial will use three packages in addition to `COGENT` – `ggplot2` and `ggthemes` for plotting and `parallel` for explicit parallelisation. You can run the script `installTutorialPackages.R` to check if you already have everything installed, and install any missing packages. All that is left to do is to load the packages and set a random seed.

```r
require("COGENT")
```

```
## Loading required package: COGENT
```

```r
require("ggplot2")
```

```
## Loading required package: ggplot2
```

```r
require("ggthemes")
```

```
## Loading required package: ggthemes
```

```r
require("parallel")
```

```
## Loading required package: parallel
```

```r
set.seed(101019)
```

This tutorial will start by describing the type of data and functions COGENT is designed to handle. You will then learn how you can use COGENT to help choose between competing network construction methods, as well as to determine suitable thresholds.

#### Gene expression data

For this tutorial we will use some yeast expression data taken from the Gene Expression Atlas. The data file `tutorialData.RData` contains a subset of the original dataset, filtered using `prepareTutorialData.R`. The data has been reduced to a set of 200 genes, expressed across 26 samples. These are stored in a data frame titled `yeastData`.

```r
load("tutorialData.RData")
head(yeastData[,1:6])
```

```
##       Name sample1 sample2 sample3 sample4 sample5
## 16     COB     0.2     0.1     0.2     0.2     0.1
## 41    SSA1    -0.3    -0.5    -0.4      NA    -0.2
## 43 YAL008W    -0.2    -0.1    -0.1     0.3      NA
## 47    CYS3    -0.2    -0.3    -0.2    -0.1    -0.3
## 48    DEP1    -0.4    -0.4    -0.5    -0.1      NA
## 55 YAL019W    -0.2    -0.2    -0.3    -0.2      NA
```

COGENT assumes a standard way of formatting gene expression data. Data should be stored in an object of class `data.frame`, in which rows represent genes and columns represent samples. Row names are allowed but will be ignored. A column titled `Name` should contain gene names. These could be set to `NA` and will not be explicitly checked. All other columns should be numeric. If you do not know whether your data is COGENT-compatible, you can check it using `checkExpressionDF()`. This will either return `TRUE` or print an error message, which should hopefully give you a hint as to what the incompatibility is.

```r
checkExpressionDF(yeastData)
```

```
## [1] TRUE
```

#### Network construction functions and network comparison

For the purposes of COGENT, a network construction function is any function which maps a COGENT-compatible data frame as above to a network adjacency matrix. The networks can be weighted, but are assumed to be undirected. The weights, if present, should be non-negative.

As mentioned earlier, one way to build a co-expression network is by calculating Pearson correlation coefficients and thresholding. Here is a simple function, which will take the top 10% of correlation values.

```
buildPearson <- function(df, quant=0.90){
  check <- checkExpressionDF(df)
  A <- cor(t(df[,colnames(df)!="Name"]), use="pairwise.complete.obs", method="pearson")
  threshold <- quantile(A[upper.tri(A)], quant, na.rm=TRUE)
  A <- 1*(A>=threshold); diag(A) <- 0
  colnames(A) <- rownames(A) <- df$Name
  return(A)
}
pearsonA <- buildPearson(yeastData)
```

Notice that rather than imposing a threshold on the raw correlation values (e.g. by insisting the correlations to be above 0.70), we fixed the top quantile of values to take, i.e. fixed the density of the obtained network. It is common to fix and control for network density, especially if later different co-expression networks are to be compared. To illustrate this, we introduce a competing way of building co-expression networks using Kendall correlations instead.

```
buildKendall<- function(df, quant=0.90){
  check <- checkExpressionDF(df)
  A <- cor(t(df[,colnames(df)!="Name"]), use="pairwise.complete.obs", method="kendall")
  threshold <- quantile(A[upper.tri(A)], quant, na.rm=TRUE)
  A <- 1*(A>=threshold); diag(A) <- 0
  colnames(A) <- rownames(A) <- df$Name
  return(A)
}
kendallA <- buildKendall(yeastData)
```

The first question to ask is whether the two networks differ in a meaningful way. The function `getEdgeSimilarity()` provides an overview of this. It takes in two adjacency matrices and compares them, returning a list of the following three elements:

- `nodeCount`, the number of nodes which have at least one edge in at least one of the networks,
- `globalSimilarity`, the (weighted) Jaccard index of the edge sets of the two networks, and
- `localSimilarity`, an array of the Jaccard indeces for each gene neighbourhood.

The purpose of `nodeCount` is to show how many nodes were used for the comparison. If input data includes many genes which have missing data or for some other reason map to isolated nodes across *both* networks, they are excluded from the analysis.

```
PKcomparison <- getEdgeSimilarity(list(pearsonA, kendallA), align=FALSE)
PKcomparison$nodeCount
```

```
## [1] 172
```

In this case, network comparison is carried over 172 genes. Some of these may have edges incident to them in only one of the two networks. The 28 genes which have no neighbours both in either the Pearson or the Kendall networks are excluded. A global similarity is then calculated in order to evaluate how similar the networks are.

```
PKcomparison$globalSimilarity
```

```
## [1] 0.6205212
```

Global similarity takes values between zero and one, with zero corresponding to no edge overlap and one corresponding to perfect overlap. Suppose we had two random networks, each with 200 nodes and network density of 0.10. On average, their intersection would be roughly 200 edges, and their union would be roughly 3800 edges. This means we would expect a global similarity of around $200/3800 \approx 0.05$ purely by chance. The observed value of 0.62 is clearly higher than random (as we would hope!) even if the Pearson and Kendall networks differ considerably.

The global similarity is a single value measuring the agreement of the two networks. The local similarity measures the agreement across each gene neighbourhood. This can be useful if we want to know not only whether the networks are very different, but also where they differ.

```
hist(PKcomparison$localSimilarity,
     main="Local similarity between Pearson and Kendall networks",
     xlab="Similarity",
     breaks=seq(0, 1, .05), col="cornflowerblue")
```

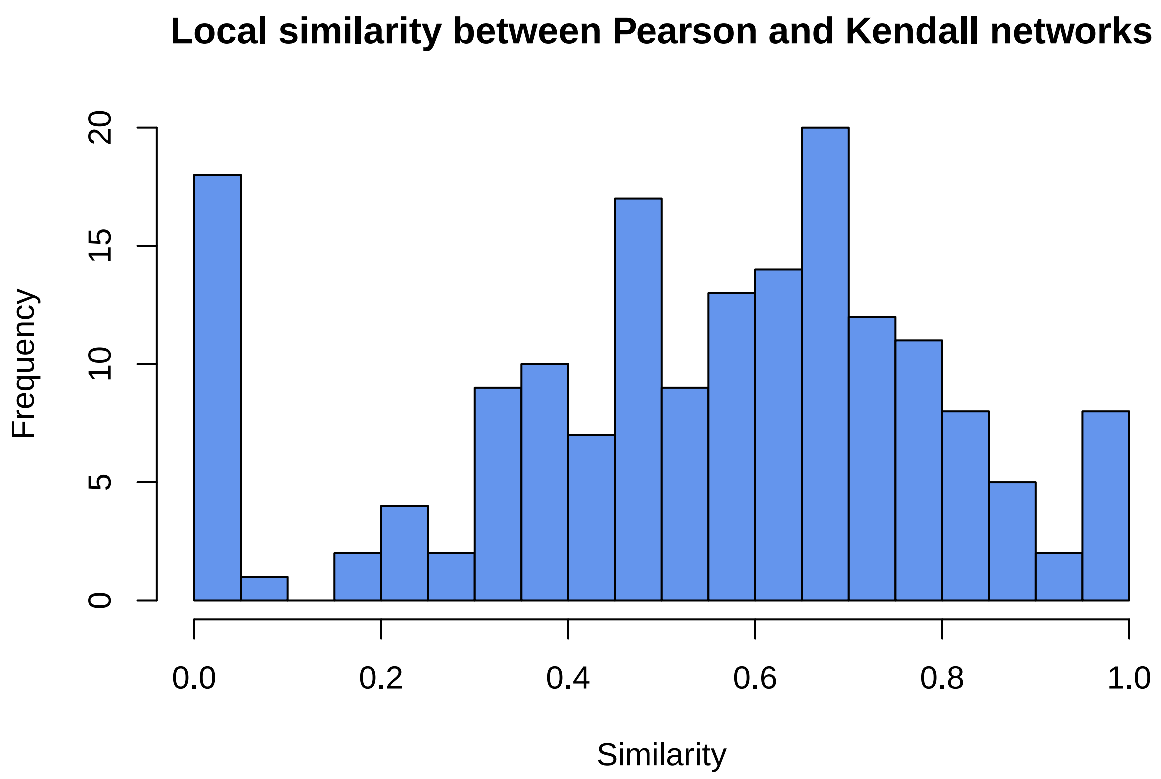


For the local similarity between these two networks, a set of low-degree nodes accounts for the peaks at 0 and 1. Even disregarding these nodes, the local similarity distribution is still rather broad (mean 0.58 and standard deviation 0.18 after removing extreme values). If we had a gene set of interest, we could inspect it in more detail at this stage.

Our initial inspection shows that while the Pearson and Kendall networks overlap, their differences are far from non-negligible. A natural question to then ask is which of the two should be prioritised for downstream analysis.

#### COGENT analysis

COGENT evaluates network consistency by repeatedly splitting the set of samples of gene expression data in two, and building a separate network from each sample subset. The more similar the resulting networks are, the more consistent the network construction method is judged to be. To measure network similarity, COGENT uses `getEdgeSimilarity()`, as well as an optional sister function, `getNodeSimilarity()`. The latter measures similarity based on a node metric – e.g. you can compare degree sequences, or betweenness centrality across two networks. The comparison itself can be done by rank k-similarity, by a correlation coefficient, or by Euclidean distance.

First, we define a node metric function to use. In this example, we use the degree. Note that the node metric function should map from an adjacency matrix to an array.

```
calculateDegrees <- function(A){
  deg <- rowSums(A)
  names(deg) <- colnames(A)
  return(deg)
}
```

The following single run of the COGENT function will split original data samples in two overlapping subsets and construct a network using `buildPearson()` from each one. The parameter `propShared` fixes the proportion of the 26 samples to be shared across the two subsets. The remaining samples will be split equally among the subsets. To ensure parity, a sample may be omitted at random. In this case, `propShared=0.50`, so that each subset will consist of 19 samples – 13 shared, and 6 unique to each dataset.

```
x <- cogentSingle(df=yeastData, netwkFun=buildPearson, propShared=0.50)
x$globalSimilarity
```

```
## [1] 0.4917541
```

We can see that subsetting the data results in networks that are more different to each other than the Pearson and Kendall networks are over the full dataset! The Kendall network builder performs similarly. We demonstrate this below, and further include node metric comparison using the `calculateDegrees()` function. After the data is randomly split and two networks are constructed, COGENT will calculate global and local similarity as before. In addition, the degree sequences of the two networks will be calculated and compared. The degree sequence comparison will be done in all possible modes – via correlation (with parameters given by `use` and `method`), via rank k-similarity (with parameter given by `k.or.p`), and via Euclidean distance. The distance can be scaled, using `scale=TRUE`. See `?getNodeSimilarity` for further details.

```
x <- cogentSingle(df=yeastData, netwkFun=buildKendall, propShared=0.50,
                  nodeFun=calculateDegrees, nodeModes="all",
                  use="pairwise.complete.obs", method="pearson",
                  k.or.p=0.10,
                  scale=TRUE)
c("globalSimilarity"=x$globalSimilarity, "corDegrees"=x$corSimilarity)
```

```
## globalSimilarity       corDegrees
##        0.5660897        0.9280868
```

Notice the difference between the global similarity (0.56) and the degree similarity, measured by Pearson's correlation coefficients (0.93). While the generated networks are considerably different, their degree sequences are very similar. If subsequent analysis will be based on the degrees, a relatively low global similarity may be unimportant.

The values above rely on random sampling. In order to understand better the consistency of each method, we can run the COGENT function multiple times and study the distribution of obtained similarities. The function `cogentLinear()` does this and aggregates the results into a data frame.

```r
stabilityPearson <- cogentLinear(df=yeastData, netwkFun=buildPearson, propShared=0.50,
                                repCount=100, nodeFun=calculateDegrees, nodeModes="all",
                                use="pairwise.complete.obs", method="pearson",
                                k.or.p=0.10, scale=TRUE)
head(stabilityPearson)
```

```
##   nodeCount globalSimilarity localSimilarity corSimilarity ksimSimilarity
## 1       185        0.5474339       0.4571917     0.9022193           0.70
## 2       183        0.4664702       0.3550761     0.8619905           0.70
## 3       178        0.4951165       0.4318332     0.8831056           0.65
## 4       180        0.7081545       0.6353801     0.9587328           0.70
## 5       182        0.6392092       0.5251718     0.9503151           0.75
## 6       181        0.3800277       0.3035073     0.7237742           0.50
##   L2Distance
## 1 0.12218187
## 2 0.15415357
## 3 0.14172801
## 4 0.07951689
## 5 0.09309951
## 6 0.21778900
```

Pearson correlations are not computationally demanding, so the above only takes a couple of seconds. However, Kendall correlations can take much longer to compute, especially with a larger set of genes. To speed the calculation up, we can parallelise using `cogentParallel()`. This can be particularly useful if your network construction method is slow but not itself easily parallelisable.

*Note: Since Windows does not support forking, you may need to set `threadCount=1` in the function call below. In this case the code will require longer to run.*

```r
stabilityKendall <- cogentParallel(df=yeastData, netwkFun=buildKendall, propShared=0.50,
                                  repCount=100, threadCount=4, nodeFun=calculateDegrees,
                                  nodeModes="all", use="pairwise.complete.obs",
                                  method="pearson", k.or.p=0.10, scale=TRUE)
head(stabilityKendall)
```

```
##   nodeCount globalSimilarity localSimilarity corSimilarity ksimSimilarity
## 1       179        0.5212075       0.4251870     0.8968095           0.55
## 2       177        0.5282150       0.4265865     0.9050937           0.65
## 3       182        0.3838665       0.2948270     0.7904382           0.50
## 4       182        0.5196640       0.3989072     0.9003559           0.65
## 5       181        0.4504734       0.3497387     0.8519830           0.55
## 6       181        0.4956783       0.4098572     0.8913320           0.65
##   L2Distance
## 1  0.1230002
## 2  0.1256727
## 3  0.1724491
## 4  0.1271666
## 5  0.1557248
## 6  0.1369928
```

Similarities obtained by subsampling in this way provide a measure of *consistency* of the network building methods. We can choose between the two types of correlation coefficients, by seeing which one is more consistent. Below we plot the global similarities obtained by the two COGENT runs.

```r
stabilityBoth <- rbind(
  cbind(stabilityPearson, method="Pearson"),
```

```r
  cbind(stabilityKendall, method="Kendall")
)
ggplot(stabilityBoth, aes(x=method, y=globalSimilarity)) +
  theme_economist_white() +
  geom_boxplot() +
  ggtitle("Stability of Pearson and Kendall\ncorrelation networks") +
  scale_x_discrete("Method") +
  scale_y_continuous("Global similarity")
```

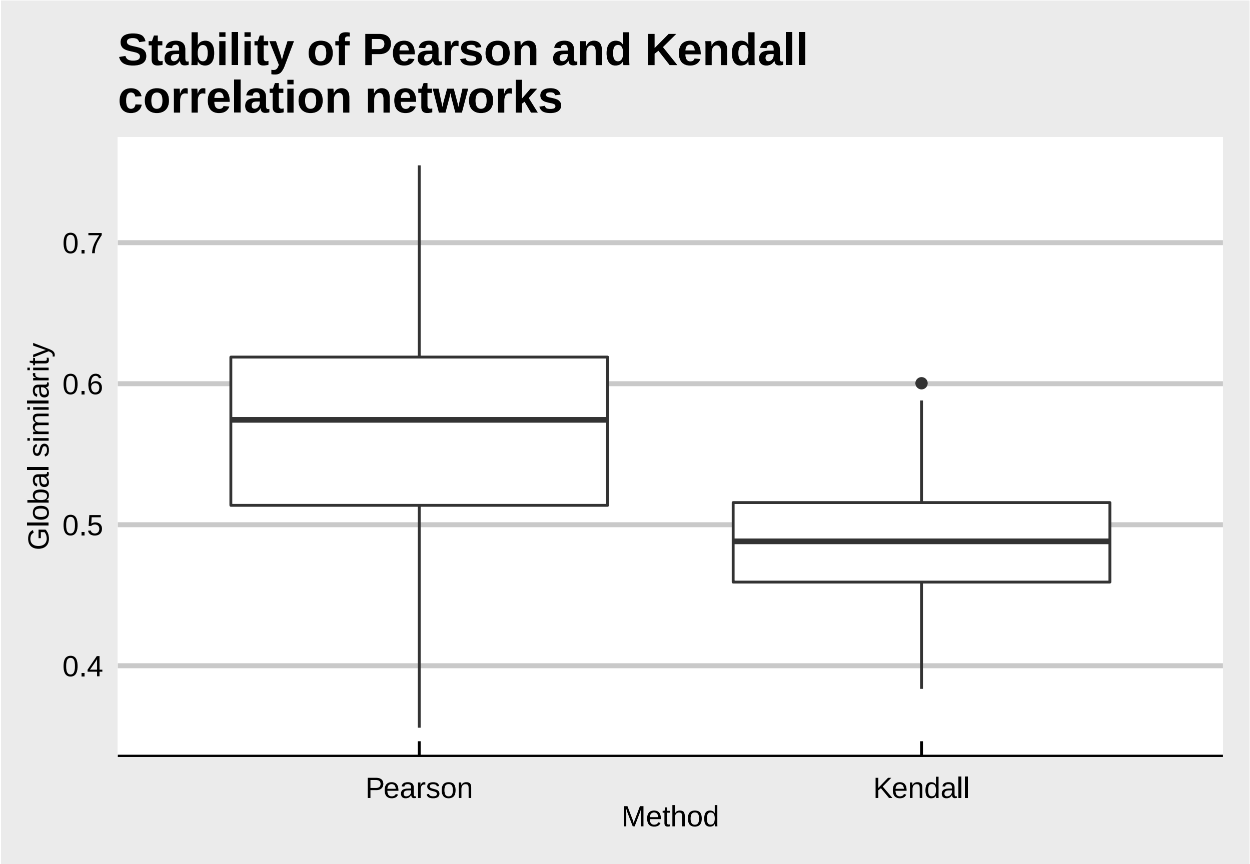


While our initial calculations showed the Kendall network builder to be a little more consistent, repeated sampling reveals the Pearson builder actually has higher global similarity. A formal test, such as a Wilcoxon rank-sum test can show whether the difference is significant. In this case, however, the picture speaks for itself.

Above we saw that the degree sequences are potentially preserved much better than the edge sets. We can also compare the two builders through the degree sequence similarity.

```r
ggplot(stabilityBoth, aes(x=method, y=corSimilarity)) +
  theme_economist_white() +
  geom_boxplot() +
  ggtitle("Stability of Pearson and Kendall\ncorrelation networks") +
  scale_x_discrete("Method") +
  scale_y_continuous("Degree consistency via correlations")
```

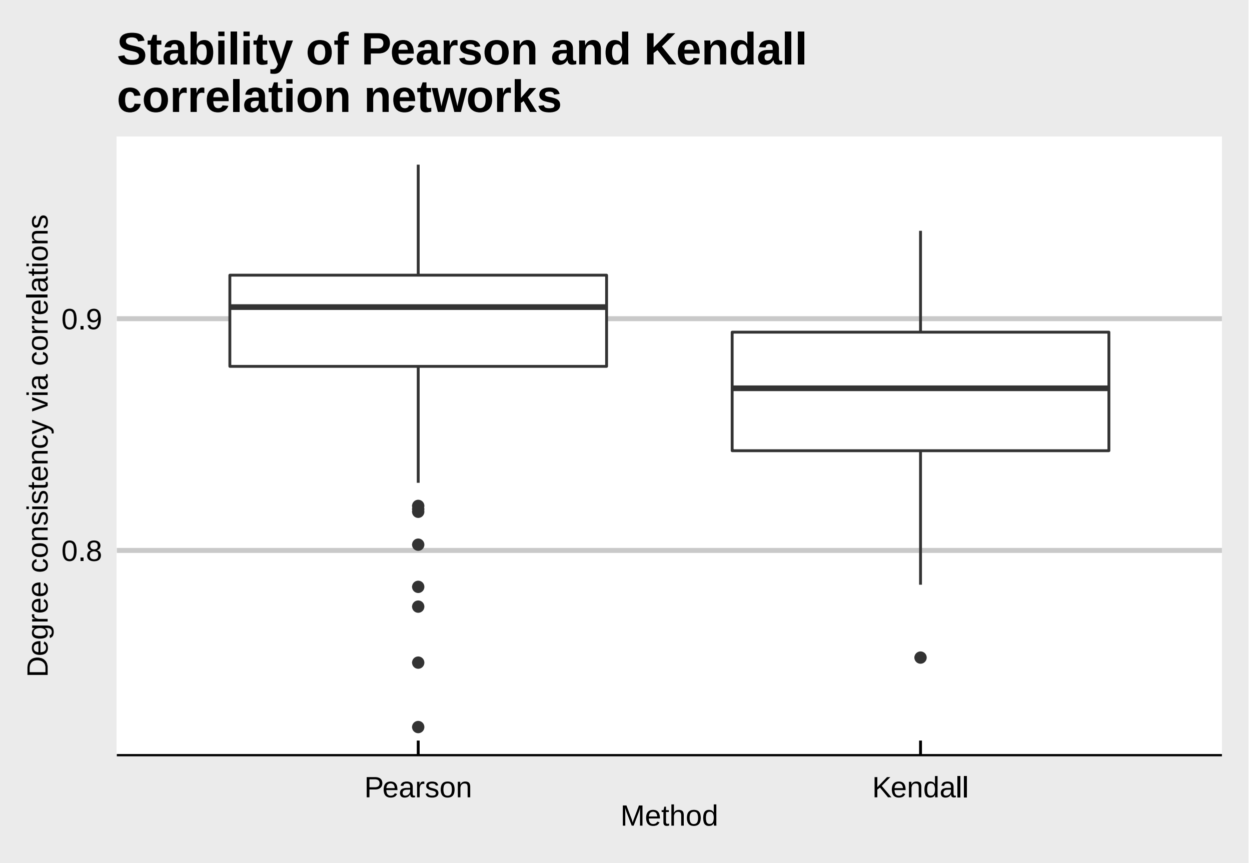


There are two things to notice on this plot. One is that both methods produce overwhelmingly similar degree sequences under subsampling (values are around 0.90). The other is that the Pearson network still outperforms the Kendall one, even if less convincingly.

Overall, we have found that the Pearson correlation network construction method performs slightly better in terms of consistency (measured both as global similarity and in terms of degree sequences) than the Kendall correlation one **for this data set**. But when we first started, we imposed an arbitrary threshold for the network density. We can also use COGENT to help inform treshold choice.

#### Choosing a threshold

Edge similarity metrics as used in COGENT will scale with the number of edges in the network: intuitively, more edges means more of them will overlap just by chance. We would expect COGENT global and local similarities to be higher for denser networks, which means they would not be very useful for setting thresholds. Consistency via node comparison is one way of circumventing this issue. Another is to correct the edge similarity metrics for network density. We do this by comparing to random networks generated with the configuration model, using `getEdgeSimilarityCorrected()`. There are two types of comparison: random and expected. The former generates configuration model networks and uses them for the correction. The latter only permutes its input degree sequences and works with approximate expectations.

The following function will take in a threshold, split the data, compute the corresponding pair of thresholded Pearson correlation networks, and compare them:

```
getThresholdStability <- function(th){
  dfSplit <- splitExpressionData(yeastData, propShared=0)
  A <- lapply(dfSplit, function(df) buildPearson(df, th))
```

```
  return(getEdgeSimilarityCorrected(A, type="expected"))
}
```

Again, we can apply it multiple times, say 100 each, at several thresholds. Here we parallelise explicitly. We threshold for network density between 0.01 and 0.50 at 0.01 intervals. This corresponds to quantiles of the correlation distribution between 0.50 and 0.99.

*Note: Since Windows does not support forking, you may need to set* `mc.cores=1` *in the function call below. In this case the code will require longer to run. If you are short on time, you can reduce the* `thresholds` *array, e.g. by increasing the gap by setting* `thresholds <- seq(0.5, 0.99, 0.05)` *instead. You can also cheat and load the output directly (see end of tutorial).*

```
aggregateThresholdStability <- function(th, repCount=100){
  thS <- replicate(repCount, getThresholdStability(th), simplify=FALSE)
  thS <- do.call("rbind", thS); thS <- apply(thS, 2, unlist)
  return(as.data.frame(cbind(thS, threshold=th)))
}
thresholds <- seq(0.5, 0.99, 0.01)
thresholdComparisonDF <- mclapply(thresholds, aggregateThresholdStability, mc.cores=6)
thresholdComparisonDF <- do.call("rbind", thresholdComparisonDF)
```

Note that in this case it is quick to split the data and generate new correlation networks at every threshold. If the calculation is slower (e.g. if we use Kendall correlations instead), we may want to do the split 100 times overall, and then simply threshold the same scored networks at the desired values.

We can now see how similarity changes with the threshold. We plot the network density, which is calculated as 1-threshold, versus the density-adjusted global similarity.

```
thresholdComparisonDF <- subset(thresholdComparisonDF,
                                !is.na(thresholdComparisonDF$correctedSimilarity))
ggplot(thresholdComparisonDF, aes(x=1-threshold, y=correctedSimilarity)) +
  geom_smooth() +
  theme_economist_white() +
  ggtitle("Threshold choice for Pearson correlation networks") +
  scale_y_continuous("Density-adjusted consistency") +
  scale_x_continuous("Network density", breaks=seq(0, 0.5, .05))
```

```
## `geom_smooth()` using method = 'gam' and formula 'y ~ s(x, bs = "cs")'
```

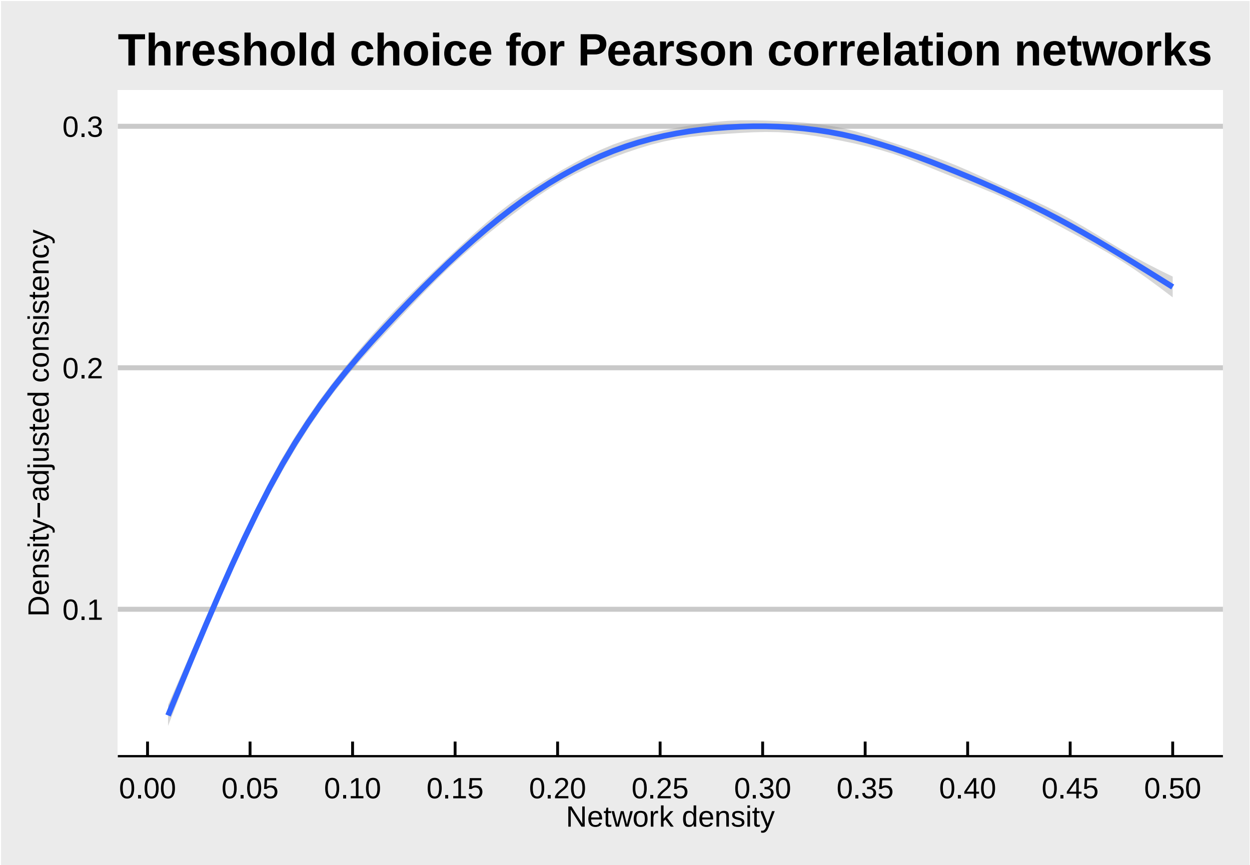


We performed our original analysis at network density of 0.10, while a network of around 0.30 would have significantly higher consistency. We could apply the same idea to the Kendall builder, and check whether there are any thresholds for which it outperforms the Pearson one. This way, a network construction method can chosen with respect to both the co-expression measure and the density threshold employed.

##### Wrapping up

Above we have demonstrated two of the main applications of COGENT: choosing between different network building approaches and determining a threshold for the construction of gene co-expression networks. We showed that, for a toy data set, networks obtained from Pearson correlations outperform networks from Kendall correlations. Further, we demonstrated that a density of around 0.30 results in the most consistent Pearson correlation networks for this dataset. If you would like to inspect the data further without generating it fresh, you can find the complete numerical output of this tutorial in the `tutorialOutputData.RData` file.

In addition to gene expression data, COGENT can be readily applied to other types of data, where co-occurrence signifies a functional relationship of interest. This could, for example, include microbiote data, as well as discrete data, such as synthetic lethality screens.
